# Tuberin levels during cellular differentiation in brain development

**DOI:** 10.1101/2021.10.17.464725

**Authors:** Bashaer Abu Khatir, Gordon Omar Davis, Mariam Sameem, Rutu Patel, Jackie Fong, Dorota Lubanska, Elizabeth Fidalgo da Silva, Lisa A. Porter

## Abstract

Tuberin is a member of a large protein complex, Tuberous Sclerosis Complex, and acts as a sensor for nutrient status regulating protein synthesis and cell cycle progression. Mutations in the Tuberin gene, TSC2, lead to the formation of tumors and developmental defects in many organ systems, including the central nervous system. Tuberin is expressed in the brain throughout development and levels of Tuberin have been found to decrease during neuronal differentiation in cell lines *in vitro*. Our current work investigates the levels of Tuberin at two stages of embryonic development *in vivo*, and we study the mRNA and protein levels during a time course using immortalized cell lines *in vitro*. Our results show that Tuberin levels remain stable in the olfactory bulb but decrease in the Purkinje cell layer during embryonic mouse brain development. We show here that Tuberin levels are higher when cells are cultured as neurospheres, and knockdown of Tuberin results in a reduction in the number of neurospheres. These data provide support for the hypothesis that Tuberin is an important regulator of stemness and the reduction of Tuberin levels might support functional differentiation in the central nervous system. Understanding how Tuberin expression is regulated throughout neural development is essential to fully comprehend the role of this protein in several developmental and neural pathologies.

**HIGHLIGHTS:**

- Tuberin protein levels are decreased in the Purkinje cell layer in later stages of embryonic development.
- Tuberin protein and mRNA levels decrease as cells undergo neuronal differentiation.
- Downregulation of Tuberin impairs neurosphere formation.
- Tuberin is implicated in the maintenance of stemness in the developing brain.

## 1. INTRODUCTION

Tuberous Sclerosis Complex (TSC) is an autosomal-dominant disorder that can be acquired through inheritance or sporadic germ-line mutations and presents with growth of benign hamartoma tumors in the brain, kidneys, and skin. TSC prevalence affects approximately 1 in 6000 live births annually and is estimated to affect roughly 1.5 million individuals worldwide (Curatolo et al., 2008; Jones et al., 1999; Maheshwar et al., 1997; Uysal and Sahin, 2020). TSC patients harbour mutations in at least one of the tumor-suppressor proteins, Tuberin (gene - TSC2), and/or its binding partner, Hamartin (gene -TSC1).

Domains of the 180 kDa Tuberin protein are involved in regulation of a vast of cell biology events (Krymskaya, 2003). RapGAP-like domain has been demonstrated to catalyse the Rheb-GTP to Rheb-GDP hydrolysis to restrict protein synthesis and cell growth via the mTOR pathway (Li et al., 2006). Activity of the Tuberin RapGAP domain is affected by post-translational modifications occurring via nutrient signalling such as growth factors (ERK) and insulin (PI3K/Akt) and direct sensing of ATP levels (AMPK) (Inoki et al., 2003; Ma et al., 2005; Manning and Cantley, 2007). Tuberin can also directly regulate the mammalian cell cycle in both G1/S and G2/M phases (Burgstaller et al., 2009; Fidalgo da Silva et al., 2011). In G1, Tuberin functions by transporting the cyclin-dependent kinase inhibitor, p27, into the nucleus thereby enhancing the ability of p27 to inhibit cell proliferation (Burgstaller et al., 2009). In G2, Tuberin directly binds and regulates the cytoplasmic localization of the mitotic cyclin, Cyclin B1 (Fidalgo da Silva et al., 2011). In high nutrient conditions, Tuberin delays nuclear import of Cyclin B1 and permits an increase in cell size, this is reversed in low nutrient conditions where Tuberin aids in rapid nuclear accumulation of Cyclin B1 (Fidalgo da Silva et al., 2011; Fidalgo da Silva et al., 2019).

Mutations in TSC2 can result in severe hamartomas of the central nervous system (CNS) that form as early as 20 weeks of gestation (Mizuguchi, 2007; Yamanouchi et al., 1997). Plethora of studies using both human tissues and rodent models, to date, have supported the potential role of Tuberin in the CNS development (Geist and Gutmann, 1995; Geist et al., 1996; Gutmann et al., 2000; Hino et al., 1993; Kobayashi et al., 1999; Li et al., 2018; Onda et al., 2002; Rennebeck et al., 1998; Yuan et al., 2012). Tuberin expression levels have been found to be elevated in the human adult brain when compared with other tissues, including heart and kidney (Geist and Gutmann, 1995). Microarray dataset analysis shows that the expression of TSC1 and TSC2 genes are highest in the human neo-cerebellum throughout the postnatal development (Li et al., 2018). In the rat hindbrain, elevated Tuberin gene expression levels of have been found in Embryonic Day (ED) 16 and onwards. (Geist and Gutmann, 1995). In addition, TSC2 homozygous mutations are embryonic lethal in Eker rat and TSC2 knock-out mice die at ED 10 due to failure of neural tube closure (Hino et al., 1993). Neuroepithelial progenitor cells from TSC2^-/-^ mice at ED10 display reduced functional differentiation forming giant cells with reduced expression of neuronal markers and increased mTOR activity (Onda et al., 2002). It was observed that the reduced levels of Tuberin expression in neurons correlates with the extension of TSC phenotypes using mouse models with conditional knock-out of TSC2 (Yuan et al., 2012). The above data support the hypothesis that Tuberin has an active and essential role in the embryonic stages of brain development (Hino et al., 1993; Kobayashi et al., 1999; Rennebeck et al., 1998).

Tuberin expression has been detected in adult rodents CNS in both neurons and astrocytes, and its expression was observed in the developmental brain of mouse embryos (ED13 and ED17) (Geist et al., 1996; Gutmann et al., 2000). In mice, neural progenitor cells start to differentiate into neurons at ED11 by asymmetric division, and neurogenesis at the cerebral cortex peaks at ED15 finishing around birth, 19-21 days. (Chen et al., 2017; Hartl et al., 2008; Qian et al., 2000). Neuron production for different brain regions takes place at distinct stages of prenatal and early postnatal life. Some regions (e.g., olfactory bulb) experience two neurogenesis peaks, representing the critical periods for generating unique cell classes. The Purkinje cells present a peak of neuron production early in brain development (ED12) when compared to the cerebral cortex and between ED13 and ED17 Purkinje cells start to form the Purkinje cell layer (Sotelo and Rossi, 2013). The different peaks of neurogenesis in brain regions shows a fine-tune regulation of neuron production during development (Chen et al., 2017) (Fig. 1).

**Fig. 1.**
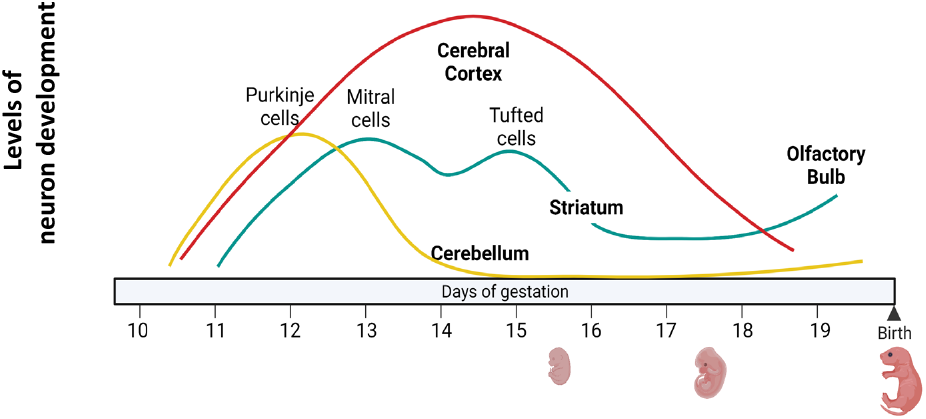
Chronology of embryonic neuron production in the developing mouse CNS. Comparison of the levels of neurons production between CNS regions of mice brain from ED10 to the time of birth. Figure adapted from (Chen et al., 2017). Image created with BioRender.com.

While an *in vivo* role for Tuberin in neural cell fate has not been conclusively addressed, data are mounting to support the potential for Tuberin in regulating neural precursor decisions. Knockdown and/or mutation of Tuberin prevents proper axon formation in developing neurons and there has been a reported increase in the proliferation of neural precursors in the cerebellum (Bhatia et al., 2009; Floricel et al., 2007; Wildonger et al., 2008). Cerebellar lesions associated to more severe clinical and neuroradiological phenotypes have been associated with TSC2 mutations (Toldo et al., 2019) and although, the cellular origin of TSC hamartomas is not known and likely variable, they are often found to express markers of immature neural stem cells, such as Nestin and Vimentin, indicating a developmental or stem cell association (Jones et al., 1999). Studies *in vitro* using PC12h (sub-clone of PC12) cells show that Tuberin levels decrease during NGF-induced neuronal differentiation (Floricel et al., 2007). Meanwhile, NOTCH mediate repression of Tuberin regulates cellular differentiation in of intestinal stem cell lineage in Drosophila (Kapuria et al., 2012). Tuberin also regulates differentiation through NOTCH in a human angiomyolipoma-derived cell line carrying a biallelic TSC2 mutation and TSC2-null rat cells. (Karbowniczek et al., 2010).

Available data support the hypothesis that Tuberin plays an important role in cell fate decisions in the developing brain. While the activity and function of the Tuberin at the cellular level are dependent on its post-translational modifications, interactions with binding partners and subcellular localization; delineating the gene and protein expression levels of Tuberin along the CNS development, throughout neurogenesis and during in cell fate determination are pivotal to understanding its regulation in the state of disease.

## 2. MATERIAL AND METHODS

### 2.1. Cell Culture

SH-SY5Y human neuroblastoma cells were obtained from ATCC. Cells were cultured in DMEM (Sigma-Aldrich) supplemented with 10% FBS (Sigma-Aldrich) and 0.25 µg/mL of penicillin/streptomycin (Gibco). DAOY human medulloblastoma cell line was obtained from ATCC and was cultured in EMEM (VWR) supplemented with 10% FBS (Gibco) and 1% penicillin and streptomycin.

### 2.2. BALB/c Neural Tissue and Primary Cell Extraction

BALB/c mice were purchased from Charles River Laboratories (Strain Code: 028). All experiments performed on animals were according to the protocols approved by the Animal Care Committee under the University of Windsor Animal Care Guidelines (AUPP# 12-18). Tissue from whole brain cerebral cortex, olfactory bulb and cerebellum were harvested from the mice. All collected tissues were flash frozen via immersion in liquid nitrogen and subsequently stored at -80°C.

The cerebellum tissue of post-natal (PN) day 4 mice was dissected to obtain primary cerebellum cells as previously described by (Pacey et al., 2006). Briefly, a midline incision was made on the skin and the skull was peeled off with sharp-pointed forceps. The spinal cord and blood vessels were cut. The brain was removed, and the cerebellum portion was placed in a 60 mm dish with DMEM and 3% penicillin and streptomycin (Gibco) and incubated on a shaker for 2 min. The cerebellum tissue was chopped with a scalpel blade and triturated with DMEM twice.

### 2.3. Immunohistochemistry Staining and Analysis

Collected tissue samples were immersed in 4% paraformaldehyde for 2 days, then a 20% sucrose bath overnight and a 30% sucrose bath overnight. Tissue was embedded in Shandon M-1 Embedding Matrix (Thermo Fischer) and sectioned at 30 µm thickness using the cryostat Leica cm3050s.

Tissue was stained by immersing the slides in 0.3% hydrogen peroxide for 30 min, washed with PBS for 5 min, then in diluted blocking serum for 20 min. Sections were incubated with rabbit anti-Tuberin polyclonal antibody (1:500) at 26° C for 1 hr or at 4° C overnight. Secondary antibody (1:500) was incubated for 1 hr at room temperature. Further staining was completed with 3,3-Diaminobenzidine substrate solution (DAB, Vector Laboratories). Sections were dehydrated through increasing concentrations of ethanol washes and cleared using xylene. Slides were mounted using Permount solution.

Tissue samples were analysed using ImageJ software. Images were opened using white to black step ladder and converted to a 32-bit image type. Analysed regions were outlined using the free-hand tool and quantified according to the mean grey value, defined as the average intensity within the region divided by the number of pixels in that region. The mean grey value of the control slides with no primary antibody was subtracted from the experimental samples to normalize the values.

### 2.4. Lentiviral Preparation and Infection

pLKO.1-TSC2 was a gift from Do-Hyung Kim (Addgene plasmid # 15478) and pLKO.1 puro was a gift from Bob Weinberg (Addgene plasmid # 8453). Lentivirus was generated as described previously (Ferraiuolo et al., 2017). Using a MOI of 10, DAOY cells were infected at 80% of confluency in a 96-well plate. 1 µL of virus was added per well with 10 µg/mL of polybrene (Sigma) in 100 µL of EMEM without antibiotics or serum overnight. The following day, cells were treated with 2 µg/mL of puromycin (Sigma) for 24 hours and allowed to recover for 24 – 48 hours.

### 2.5. Induction of Neuronal Differentiation

Primary cerebellum cells were maintained in serum free neurobasal media supplemented with B27, 20 ng/mL EGF (Gibco; #12483) and 10ng/mL EGF (Sigma; #1540). The cells were seeded at 5×10^4^ cells per well in an ultra-low attachment 6-well plate (Corning; #3471). Cells were incubated at 37° C in 5% CO_2_. Every 5-10 days, the primary neurospheres were dissociated using 1 mL 0.05% Trypsin (HyClone; #SH3023601). Once the sub-cultured primary sphere cells were needed for differentiation assays, the cells were cultured with media supplemented with 2% Fetal Bovine Serum (FBS) (Gibco; #12483). Fresh media was added to the plate every 2-3 days to enrich the most stem-like population of cells.

SH-SY5Y cells at 60 – 70% confluency were induced to differentiate using 13-cis retinoic acid at a concentration of 2 µM (Sigma-Aldrich) in the growth media. The inducing media was renewed every two days. Full neuronal differentiation was confirmed by the development of extended neurite processes measuring at least 2x the diameter of the cell body (Simpson et al., 2001) .

### 2.6. Western Blot

For western blot analysis, cells were lysed with a 0.1% NP40 or 1% Triton lysis buffer containing 0.05mM EDTA, 0.1 M NaCl, 0.02 M Tris pH 7.5, 5 µg/mL aprotinin, 0.1 mg/mL leupeptin, and 5 µg/mL PMSF. 20 ug to 200 ug of protein was run on a 7.5 or 10% SDS-PAGE gel and transferred onto a PVDF membrane (Sigma-Millipore). The membranes were blotted for 1 hr in 1, 2 or 5 % milk or BSA in TBST solution. After blocking, membranes were incubated with primary antibodies at 4° C overnight, then incubated for 1 hr in HRP-conjugated secondary antibody and washed in TBST. Membranes were incubated using the Pierce ECL Substrate (Thermo) and imaged using the FluorChem 9000 Imaging System (Alpha Innotech) or BioRad imager. Primary antibodies used in western blot analysis were: Tuberin (XP) (1:1000; Cell Signalling 4308), Tuberin (1:1000; Cell Signalling 3612), Tuberin (C20) (1:500; Santa Cruz sc-893), and Actin (1:1000; Millipore MAB1501R). Secondary antibodies used were anti-mouse IgG-peroxidase (Sigma-Aldrich Co. – Product #: A9917), and anti-rabbit IgG-peroxidase (Santa Cruz - Product #: sc-2020).

### 2.7. Neurosphere Formation Assay

SH-SY5Y neurosphere formation assays were performed as described by Pacey et al. (2006) to isolate and enrich the most stem-like cells within cultured SH-SY5Y cell populations. Briefly, 5×10^4^ cells were seeded onto ultra-low adherence 6-well culture plates in 2 mL of serum-free DMEM-F12 media (Sigma-Aldrich), supplemented with 10 µg/mL Putrescine, 5 mM HEPES, 5 µg/mL Insulin, 5 µg/mL Transferrin, 0.012% Glucose, 0.125 µg/mL Sodium Selenite, 6.3 ng/mL Progesterone and incubated for 2 to 7 days to allow for sphere formation.

DAOY cells were sub-cultured onto low adherent 6-well plates (Greiner Bio-One) with neurobasal media containing 1X B27 supplement, 12.5 ng/mL of fibroblast growth factor (FGF), and 20 ng/mL of epidermal growth factor (EGF). Upon formation of primary neurospheres, they were dissociated with 0.1 mM EDTA to and replated to form secondary neurospheres. The number of neurospheres was scored per the total number of spheres formed in each well.

### 2.8. RNA Isolation and Quantitative Real-Time PCR (qRT-PCR)

Total RNA was extracted from cells and tissues using the RNeasy Plus Mini Kit (Qiagen). cDNA was synthesized using Superscript II Reverse Transcriptase (Invitrogen). Gene expression levels were assessed using the SYBR-Green qPCR method. qRT-PCR was performed in 96-Well Clear Half-Skirt PCR Microplates (Axygen). The PCR reaction had 10 µl of SYBR-Green Master Mix (SA Biosciences Inc), 2 µL of cDNA, 1 µL of a 10 µM gene specific forward and reverse primer mixture, and nuclease free water to have a final volume of 20 µL. The primers designed for these assays were for human and mouse variants.: human TSC2 (For: 5’-GAGAGGAGCCGTGTTTTTTGTG-3’/ Rev: 5’-GACATGCCATGGCCTGGTA-3’), mouse TSC2 (For: 5’-TTGTGAGGAGGTTATGGCCATT-3’/Rev: 5’-GCAGCTGAACTTCATCTCTGTTGTAG-3’), mouse Nestin (For: 5’-ACCTATGTCTGAGGCTCCCTATCCTA-3’/ Rev: 5’-ACCTATGTCTGAGGCTCCCTATCCTA-3’) and human Oct 4 (For: 5’-CTTGCTGCAGAAGTGGGTGGAGGAA-3’/Rev: 5’-CTGCAGTGTGGGTTTCGGGCA-3’) PCRs were conducted using an ABI 7300 Real Time PCR System (Applied Biosystems). PCR cycling conditions were 95° C for 10 min.; 40-60 cycles: 95° C for 15 sec., 60° C for 1 min. mRNA levels were analysed through relative quantification study using the ABI 7300 System Sequence Detection Software v1.3 (Applied Biosystems). Analysis of obtained CT values and mRNA quantification were conducted using the comparative ddCT method, with values represented as the mean log10 RQ ± standard error (S.E.). Data analysis was conducted using Microsoft Excel (Microsoft).

### 2.9. Statistical Analysis

Evaluation of western blot data was conducted using the log 10 transformed densitometry data. Statistical analysis of changes between individual time points was conducted using Student’s T-test. Two-way ANOVA was also used to compare protein level changes, across full time courses, between selected markers. Linear regression analysis, comparing protein level changes to time, was used to evaluate trend changes across each differentiation/developmental time course. Spearman’s correlation analysis was used to compare time course trends between proteins.

## 3. RESULTS

### 3.1. Tuberin levels are regulated during the development of the CNS

The levels of Tuberin were investigated in the cerebral cortex, olfactory bulb, and Purkinje cell layer via immunohistochemistry across embryonic developmental timepoints. Six to nine sagittal mouse brain sections per timepoint over up to 4 mice were stained and quantified to conduct comparative analysis of Tuberin expression levels in select CNS regions at ED14.5, a mid-point of peak neurogenesis, and at ED16.5 just before the start of gliogenesis (Qian et al., 2000) (Fig. 2).

**Fig. 2.**
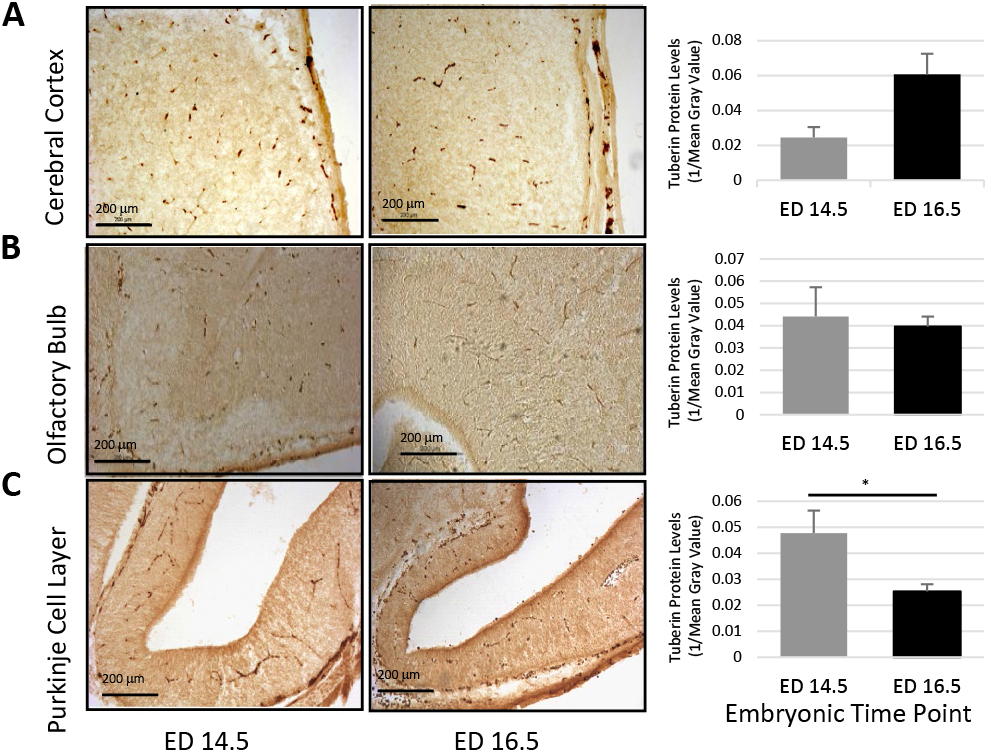
Tuberin levels in the CNS during embryonic development. Sagittal brain sections of the mouse brain obtained at ED14.5 and ED16.5 were incubated with polyclonal Tuberin antibody and DAB stained, then quantified according to the mean grey value, the average intensity within the region divided by the number of pixels in that region. Between six to nine sections from 3 to 4 mice were examined for each day. Statistical analysis was done using Anova single factor and the P values are 0.07, 0.87 and 0.01 for A, B, C respectively.

No statistical difference was found when comparing time points for the cerebral cortex (Fig. 2A) and olfactory bulb (Fig. 2B). The levels of Tuberin in the cerebral cortex, although not statistically significant, increase at ED16.5 when compared to ED14.5. This trend correlates with the decrease in the production of neurons demonstrated by Chen et al. (Chen et al., 2017) and depicted in Fig. 1.

We observed a significant decrease in Tuberin protein levels in the PCL at E16.5 when compared to E14.5 (Fig. 2C). As previously reported, Tuberin plays an important role during axonogenesis (Bhatia et al., 2009; Floricel et al., 2007; Wildonger et al., 2008), and the higher protein levels of Tuberin at ED14.5 compared to ED16.5 in the PCL suggest the role of Tuberin in the earlier stages of axon formation. Our results support that Tuberin levels are constitutively expressed throughout the CNS and may play an important role in select regions during the embryonic development.

### 3.2. The decreasing of Tuberin levels stimulates the differentiation of the stem cell population

Granular cells in the cerebellum present intense neurogenesis between ED18.5 and PN16 of the mouse brain development (Chen et al., 2017; Corrales et al., 2006; Sudarov and Joyner, 2007). To quantify the mRNA levels of TSC2 during differentiation of cerebellar progenitors, we established primary cell cultures from cerebella obtained from mice at PN day 4. The cells were subjected to differentiation *in vitro* by culturing in low-adherent plates in neurobasal media, for 72 hours and TSC2 mRNA levels were quantified via qRT-PCR. Nestin, stemness marker, expression levels were analysed to measure changes within the stem and/or progenitor cell population (Wiese et al., 2004). At 72 hours, the mRNA levels of both, TSC2 and Nestin, demonstrated a decrease of approximately 50% when compared to non-differentiated control (Fig. 3A).

**Fig. 3.**
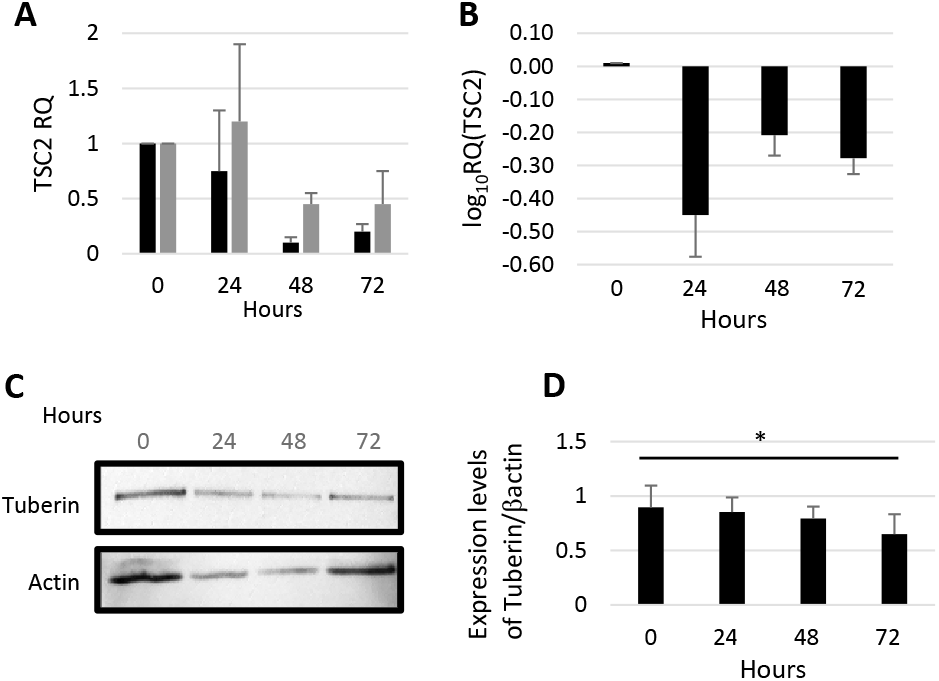
Stem cell differentiation is regulated by Tuberin levels in the CNS. (A) mRNA expression analysis of primary cerebellum cells. Post-natal day 4 cells were sub-cultured into primary neurospheres, dissociated, and differentiated over 72 hours by supplementing serum-free neurobasal medium with 2% FBS. TSC2 (black bars) and Nestin (grey bars) levels normalized to GAPDH and reported as relative quantification value (RQ). Error bars indicate SEM of three independent qrt-PCR experiments ran in triplicate. mRNA levels are normalized against 0 time point. p> 0.05. Statistical analysis was done one-way ANOVA. (B) SH-SY5Y cells were subjected to 2 µM of 13-cis retinoic acid and TSC2 RNA levels were measured for 72 hours. (C) Tuberin protein levels measured during the differentiation of SH-SY5Y cells and quantified as relative Tuberin expression to βactin. Anova statistical test *p<0.05.

Additional assessment of the Tuberin levels during differentiation was conducted using an immortalized human SH-SY5Y neuroblastoma cell line, commonly used as a model of neuronal differentiation (Shipley et al., 2016). Cells were stimulated to undergo differentiation using 2 µM of 13-cis-retinoic acid. Across the 72-hour time course, the levels of TSC2 mRNA displayed consistent downregulation at each time point tested (Fig. 3B). The regulation of TSC2 mRNA expression levels during neuronal differentiation suggest not only that dynamic variation in TSC2 expression is required for proper neuronal differentiation, but also that TSC2 expression is controlled at transcriptional level during the process.

The protein levels of Tuberin during differentiation were investigated using SH-SY5Y across a 72-hour time course (Fig. 3C). Regression analysis revealed a significant decrease of Tuberin protein levels across the time course (Fig. 3C). The obtained results support the data by Floricel *et al*., demonstrating that Tuberin protein and gene expression levels are downregulated during neuronal differentiation in PC12h cells (Floricel et al., 2007). Obtained results further support the hypothesis that Tuberin may have functional importance in maintaining stemness properties in neural populations in the brain.

### 3.3. Tuberin protein plays a functional role in regulating stemness

To determine the mRNA levels of TSC2 in stem-like populations of cells, SH-SY5Y cell line was cultured in conditions supporting self-renewal and stem cell enrichment via neurosphere formation (Fig. 4A). qRT-PCR analysis demonstrated that in comparison to cells cultured as a monolayer, there was an upregulation of OCT4 mRNA levels in the cells cultured as neurospheres, confirming their stem-like character in the applied conditions (Fig. 4A). We found the mRNA levels of TSC2 to be upregulated in SH-SY5Y neurospheres when compared to the monolayer culture (Fig. 4A). These results suggest that Tuberin has a potential role in cell populations of stem-like character. We then analysed Tuberin protein expression levels in sequential neurosphere generations from DAOY cells derived from a medulloblastoma tumor of cerebellar origin. The results demonstrated significantly elevated levels of Tuberin protein in both primary and secondary neurosphere cultures, in comparison to monolayer (Fig. 4B). To assess whether Tuberin plays a role in maintenance of self-renewing characteristics in neural cultures, DAOY cell line was manipulated to achieve Tuberin knockdown status using lentivirus and shTSC2 cells along with the scrambled control were cultured in stem-like conditions supporting neurosphere formation. The number of primary neurospheres generated by shTSC2 cells was significantly decreased compared to the control (Fig. 4C). We monitored the rate of the secondary neurospheres formation over time and found that the number of neurospheres at day 8 of culture was significantly downregulated in shTSC2 DAOY-derived spheres when compared to the number of control derived spheres (Fig. 4D). These results support that Tuberin plays a role in maintaining the stemness of select neural cell populations.

**Fig. 4.**
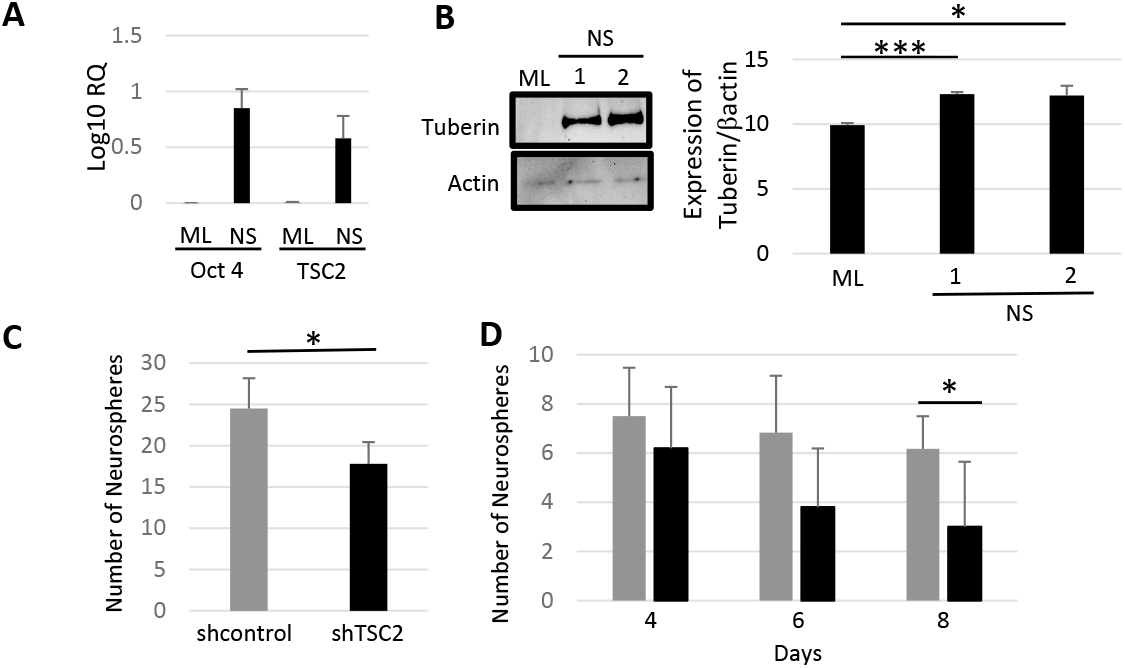
Tuberin is necessary for neurosphere formation in neuro cells. (A) TSC2 RNA levels in SH-SY5Y cells neurospheres, Oct 4 is used as stemness control. (B) Tuberin levels were determined in DAOY cells cultured in ML (monolayer), primary (1) or secondary (2) neurosphere cultures. (C&D) Comparison of the neurosphere formation in DAOY cells lines infected with shcontrol or shTSC2 to knockdown TSC2. The cells cultured in monolayer were infected with the shDNA and selected with puromycin. The surviving cells were cultured in neurosphere media and the number of primary (C) and secondary (D) neurospheres along the time course was determined. (D) shcontrol (grey bars) and shTSC2 (black bars). Anova statistical analysis *p<0.05 or ***p<0.0005.

## 4. DISCUSSION

Cell growth and proliferation are essential for normal development but must be tightly regulated to prevent tumor formation (Bai and Jiang, 2010). This represents a particular challenge during early development when cell growth and proliferation are occurring rapidly. Today, we know a great deal about specific DNA mutations and deletions that are associated with a plethora of overgrowth disorders; however, proposing suitable treatments for these disorders requires a complete understanding of how these changes impact growth and development. Mutations in the gene encoding Tuberin, TSC2, are a prominent alteration in benign tumors, known as hamartomas, which can form in many organs in the body including the brain (Zeng et al., 2011). TSC2 mutations are also found in select brain cancers such as giant cell astrocytoma (Gutmann et al., 2000). In combination with other mutations such as those in the Sonic Hedgehog pathway, Tuberin mutations enhance the prevalence of the childhood brain cancer medulloblastoma (Bhatia et al., 2009). Understanding how Tuberin regulates the growth of neural cell populations may point to therapeutic directions that can benefit patients with proliferative diseases.

The most well-established role for Tuberin is as a negative regulator of cell growth and proliferation through mTOR pathway and cell cycle progression through G1 and G2 (Burgstaller et al., 2009; Fidalgo da Silva et al., 2011; Schmelzle and Hall, 2000). mTOR is highly conserved and it is involved in a plethora of cellular processes but is most noted for controlling protein synthesis (Wullschleger et al., 2006). Hence the most obvious role for a Tuberin mutation or deletion would be to encourage cell overgrowth via a lack of control over protein synthesis and depleting the tumor suppressor roles of Tuberin on the cell cycle. These functions are likely the major drivers of several of the benign hamartomas that form in TSC2 mutated patients. Previous studies done by Kobayashi *et al*. in 1996, have shown that TSC2 knock-out mice died at ED10 due to failure of neural tube closure, suggesting the intriguing possibility that Tuberin may have an active and essential role in the early stages of brain development (Orimoto et al., 1996).

To study normal protein expression in a tissue specific manner, a Balb/C mouse model was used and IHC staining was performed in certain regions of the CNS. Tuberin protein levels were constitutively expressed across all regions of the brain studied including the olfactory bulb, cerebral cortex, and PCL. However, when protein levels within the regions were quantified between time points, it was observed that Tuberin protein levels significantly decrease in the PCL at ED16.5 when compared to ED14.5. Whether this relates to functional changes in neurogenesis within this region requires further dissection of Tuberin levels within relative cell populations through each developmental time point. It is documented in a study by Murthy *et al*. in 2001., that Tuberin expression is relatively high in the brain during brain development especially starting at ED13. Conversely, Tuberin levels were decreasing in other organs in the body including the kidneys, liver, and heart (Murthy et al., 2001). In these organs, it is possible that the cells were terminally differentiated; therefore, these cells do not need a high level of Tuberin to keep cellular proliferation and growth under control. This may shed light on the significance of Tuberin in regulating growth in mammalian brain development at early embryonic time points especially during neurogenesis. The increased protein levels of Tuberin observed in the PCL in early development may be an indication of its regulatory role in cellular growth especially in the neural stem cells. Tuberin may be allowing for cellular proliferation and preventing neural cell differentiation at this time. Analysing normal Tuberin levels within the PCL is important because neurological disorders, such as Autism Spectrum Disorders, tend to arise in the PCL of the cerebellum due to TSC2 mutations (Li et al., 2018; Reith et al., 2011). This provides early evidence for a potential role for Tuberin in this region.

Interestingly, regulation by Tuberin in the regions that give rise to the cerebellum may imply its importance in paediatric brain cancers such as medulloblastoma (Bhatia et al., 2009). Medulloblastoma arises from a subpopulation with cancer stem cell properties (Singh et al., 2004). Hence, we optimized the isolation of primary mouse cerebellum cells to study the importance of Tuberin during cell fate and differentiation decisions. At 72 hours the mRNA levels of Tuberin significantly decreased. To examine expression during differentiation, we utilized qRT-PCR to quantify TSC2 mRNA levels at each time point during differentiation. Our data showed that Tuberin mRNA is significantly decreased by 24 hours during differentiation, presenting with a biphasic pattern of expression during the first two hours, and subsequently decreasing below initial levels thereafter. The cell line used in this study are known to be comprised of heterogeneous cell populations containing stem-like progenitor cells and other, more committed precursor cells (Biagiotti et al., 2006; Neville et al., 2009). Hence, data obtained from our studies and others can be said to reflect the course of differentiation from an intermediate ‘stem-ness’ state to a terminal neuronal phenotype. This supports the hypothesis that Tuberin is needed early on before differentiation in the stem cell or progenitor population to regulate cellular growth and differentiation. The decrease in Tuberin would facilitate an increase in protein synthesis, allowing the production of specific proteins needed for fully functional differentiation. It has been previously established that Tuberin interacts with a variety of pathways that guide and regulate cell fate and differentiation. In a study of neuroepithelial progenitor cells from TSC2 homozygous null mice, cells cultured from ED10.5 had abnormal cell differentiation (Onda et al., 2002). In a separate study, Tuberin knockdown prohibited proper axon formation in actively developing neurons and an increase in proliferation of the neural precursor cells and a decrease in neuron outgrowth were reported (Floricel et al., 2007).

Tuberin expression levels were measured during *in vitro* neuronal differentiation using the human neural precursor cell line SH-SY5Y. When induced to differentiate, these cells showed downregulation of Tuberin levels. The role of Tuberin in cell differentiation has been controversial over the years. It has been observed an upregulation in Tuberin levels following neural growth factor-induced differentiation of PC12h cells, with levels remaining consistent for 72 hours and diminishing thereafter (Floricel et al., 2007), yet another study showed reduced Tuberin levels across a 30 min time course, following induction of differentiation in PC12 cells (Wu and Wong, 2005). Here, we provide results suggesting a progressive decrease of Tuberin levels during a 72 hours-time course.

Most observations of Tuberin expression and regulation have often neglected to include investigation of the potential role transcriptional may play in the regulation of Tuberin levels, due largely to a general assumption that Tuberin and many of the other TOR proteins are constitutively expressed (Miloloza et al., 2000; Soucek et al., 1997; Soucek et al., 1998). Cultured neurospheres from normal SH-SY5Y cell culture were used to measure the level of Tuberin expression in a stem-like progenitor cells comparing with the TSC2 mRNA of cell growing in monolayer under normal culture conditions. It is notable that Tuberin levels mimicked that of the stemness marker Oct4. It is possible that due to the high numbers of cells used in this assay, and the lack of clonal passaging, that we did not obtain a pure population of stem cells. Given the similarities in the expression between TSC2 and a subset of stemness markers it is certainly important to follow up these experiments using clonal passaging of a small numbers of cells. Should TSC2 mRNA levels remain stable regardless of differentiation status within these precursor cell populations, this would support the observation made by Soucek et al. (1998), indicating that TSC2 mRNA levels are indeed not subject to change because of differentiation in neuronal cells (Soucek et al., 1998).

To determine if Tuberin plays a functional role in the stem cell population, the levels of Tuberin expression were determined in DAOY cells cultured as monolayer, primary and secondary neurosphere conditions. The levels of Tuberin significantly increased when the cells are forming neurospheres. To further determine the importance of Tuberin for keeping the stem cell population, the DAOY cells were manipulated by lentiviral infection to knockdown Tuberin. Cells infected with a lentiviral shRNA scrambled vector served as the control for the experiment. There were fewer neurospheres formed when Tuberin was knocked down. This data confirms that Tuberin plays a critical role in the maintenance of the stem cell population; whether it is through self-renewal, survival or proliferation remains to be further determined. We assess self-renewal through secondary neurosphere formation in Tuberin knockdown cells. Over an 8-day culture period, there was a significant decrease in Tuberin in the secondary neurospheres. This result supports that Tuberin is essential in regulating the stem cell population.

## 5. CONCLUSION

We have demonstrated that the regulation of Tuberin expression, at the protein level, and at the mRNA level, is among the cohort of events that mark the process of neural cell differentiation as a component of the process of cell fate determination. However, there is still much that we do not know about the degree and significance of the involvement of Tuberin in this process. Further dissecting the pathways by which Tuberin may be important in regulating neural cell fate, as well as how Tuberin levels are regulated in specific cell types and tissue types are important for a better understand of cell differentiation and development. Clearly defining the roles of Tuberin during neural development may reveal new perspectives from which to examine the process of cell fate choice and new methods by which to approach the study, diagnosis, and treatment of TSC. Overall, our results show a potential role for Tuberin in regulating and maintaining the progenitor stem cell population. We have determined that Tuberin is present, and even elevated, in specific regions of the brain at early time points of development when neurogenesis is at its peak. Tuberin seems to be regulating the process of cell fate decisions in instances where it halts cell proliferation and allows for terminal cell differentiation. Assessing Tuberin levels in primary extracted cerebellum cells is one of the few studies that try to link Tuberin regulation with stem cell fate decisions. Being able to correlate Tuberin expression with tumor stem cells, such as those in medulloblastoma cells, will help imply how these cancer stem cells are driving tumorigenesis.

## Acknowledgement

We thank the Imaging and Animal facilities at the University of Windsor and Jiamila Maimaiti, Antonio Roye-Azar and Bob Hodge for technical assistance.

## Funding

This work was supported by a Seeds4Hope grant from Windsor Essex County Cancer Center Foundation (to EFS) and by a Natural Sciences and Engineering Research Council of Canada (RGPIN/06636-2014) (to LAP).

## Notes

### Competing Interest Statement

The authors have declared no competing interest.

